# Data-driven discovery of targets for bipotent anticancer drugs identifies Estrogen Related Receptor Alpha

**DOI:** 10.1101/2021.10.25.465724

**Authors:** Avinash D. Sahu, Xiaoman Wang, Phillip Munson, Jan Klomp, Xiaoqing Wang, Shengqing Gu, Gege Qian, Phillip Nicol, Zexian Zeng, Chenfei Wang, Collin Tokheim, Wubing Zhang, Jingxin Fu, Jin Wang, Nishanth U. Nair, Joost Rens, Meriem Bourajjaj, Bas Jansen, Inge Leenders, Jaap Lemmers, Mark Musters, Sanne van Zanten, Laura van Zelst, Jenny Worthington, Myles Brown, Jun S. Liu, Dejan Juric, Cliff A. Meyer, Arthur Oubrie, X. Shirley Liu, David E. Fisher, Keith T. Flaherty

## Abstract

Drugs that kill tumors through multiple mechanisms have potential for broad clinical benefits, with a reduced propensity to resistance. We developed BipotentR, a computational approach to find cancer-cell-specific regulators that simultaneously modulate tumor immunity and another oncogenic pathway. Using tumor metabolism as proof-of-principle, BipotentR identified 38 candidate immune-metabolic regulators by combining epigenomes with bulk and single-cell tumor transcriptomes from patients. Inhibition of top candidate ESRRA (Estrogen Related Receptor Alpha) killed tumors by direct effects on energy metabolism and two immune mechanisms: (i) cytokine induction, causing proinflammatory macrophage polarization (ii) antigen-presentation stimulation, recruiting CD8^+^T cells into tumors. ESRRA is activated in immune-suppressive and immunotherapy-resistant tumors of many types, suggesting broad clinical relevance. We also applied BipotentR to angiogenesis and growth-suppressor pathways, demonstrating a widely applicable approach to identify drug targets that act simultaneously through multiple mechanisms. BipotentR is publicly available at http://bipotentr.dfci.harvard.edu/.

**One-Sentence Summary:** BipotentR identifies targets for bipotent anticancer drugs, as shown by the energy and immune effects of ESRRA inhibition.

## A data-driven approach to identifying bipotent targets

Tumors frequently do not respond to drugs, while those that respond often relapse by developing drug resistance. To counteract this, treatments have been designed to simultaneously target two non-overlapping oncogenic pathways(*1*). Single drugs that are tolerated yet concurrently affect two oncogenic pathways (bipotent drugs) have the potential to provide greater clinical benefits, be effective in distinct clinical populations (*2*), and be an effective strategy against drug resistance (*3*, *4*). Few drugs are known to be bipotent, and the few known examples were discovered serendipitously. Most notable are CDK4/6 inhibitors (cell cycle inhibitor and immunomodulatory(*5*)), IMiDs (Immunomodulatory imide drugs; antiangiogenic and immunomodulatory(*6*)), and itaconate (energy metabolism and immunity(*7*)). Likewise, few bipotent gene targets have been identified, with examples including HDAC6(*8*), CDC7(*1*), and PTPN3(*9*). Bipotent gene targets may be relatively common, yet undiscovered due to the lack of systematic approaches to identify them.

We developed a data-driven approach that analyzes bulk tumor transcriptomes, single-cell transcriptomes, and chromatin accessibility data to predict genes targets that have the potential to eliminate tumors in two ways: through stimulating immune-mediated tumor elimination and suppressing a second pathway that is essential for tumor development, such as energy metabolism, angiogenesis, evasion of growth suppressor, metastases, or replicative immortality(*10*).

Tumors alter their energy metabolism to meet higher bioenergetic needs and sustain rapid proliferation (*10*). As a result, tumors often have increased dependency on oncogenic energy metabolism pathways (*11*, *12*), and targeting them metabolism can inhibit tumors (*13*, *14*). Oncogenic energy metabolism helps cancer cells to evade immune response (*15*, *16*). For example, increased glucose uptake by cancer cells (the Warburg effect) (*17*) dampens immunity in tumors (*18*) because of reduced glucose availability for effector T-cells. Past attempts to target energy metabolism have failed to show therapeutic benefits in patients (*19*–*21*) due to an inability to target energy metabolism without curtailing T-cell function. While energy metabolism is necessary for all cells, its upstream transcription factors or chromatin regulators (TFCRs) are often cell-type-specific, allowing the possibility of effects specific to cancer and immune cells (*22*, *23*). Targeting energy metabolism has the potential to kill tumors in two ways: directly through cell-intrinsic mechanisms (*24*, *25*) and indirectly through immune-mediated mechanisms (*26*). For these reasons, we chose to pair energy metabolism with immunity for initial analysis by BipotentR.

BipotentR consists of two modules, “regulation” and “immune” (Fig 1A). The regulation module predicts regulators of the input pathway(s) chosen by the user, while the “immune” module identifies immunomodulatory TFCRs. To identify bipotent regulators of energy metabolism and immune response (“immune-metabolic” regulators), we used four energy metabolism pathways with a reported role in immunity (*27*–*32*) as input for the regulation module of BipotentR: glycolysis, oxidative phosphorylation (OXPHOS), tricarboxylic acid cycle (TCA cycle), and fatty acid metabolism (FA). The regulation module estimates the affinity of ∼700 individual TFCRs to bind cis-elements near input pathway genes by mining 24,000 ChIP-seq samples (*33*, *34*). For a given TFCR, BipotentR derives its core binding sites by combining all ChIP-seq samples, and then estimates its binding affinity while controlling for sample-specific confounding effects using a linear mixed model (Methods). BipotentR identified previously known and new TFCRs (Fig 1B, Table S1). Previously known TFs included ESRRA (*35*–*37*) and its co-activator BCL3 (*38*) (both regulators of OXPHOS, glycolysis, and TCA cycle), PPARG (*39*) (an FA metabolism regulator), and CEBPB (a glycolysis regulator) (*40*). Known chromatin regulators included histone demethylase KDM5A, a regulator of OXPHOS and TCA (*41*), and histone deacetylase SIN3A, a mitochondrial gene regulator (*42*). Newly identified TFCRs included NPAT and DLX1 (OXPHOS) along with NFIC and CBX1 (TCA cycle). To ensure that ChIP-seq binding data was linked to expression and that in vitro experiments are relevant in patient samples, we asked how well the expression of predicted regulators correlated with the expression of genes in the regulated pathway across 5,000 human transcriptome datasets (*43*). We found a strong correlation for all four energy metabolism pathways (Fig S1B-E).

**Figure 1.**
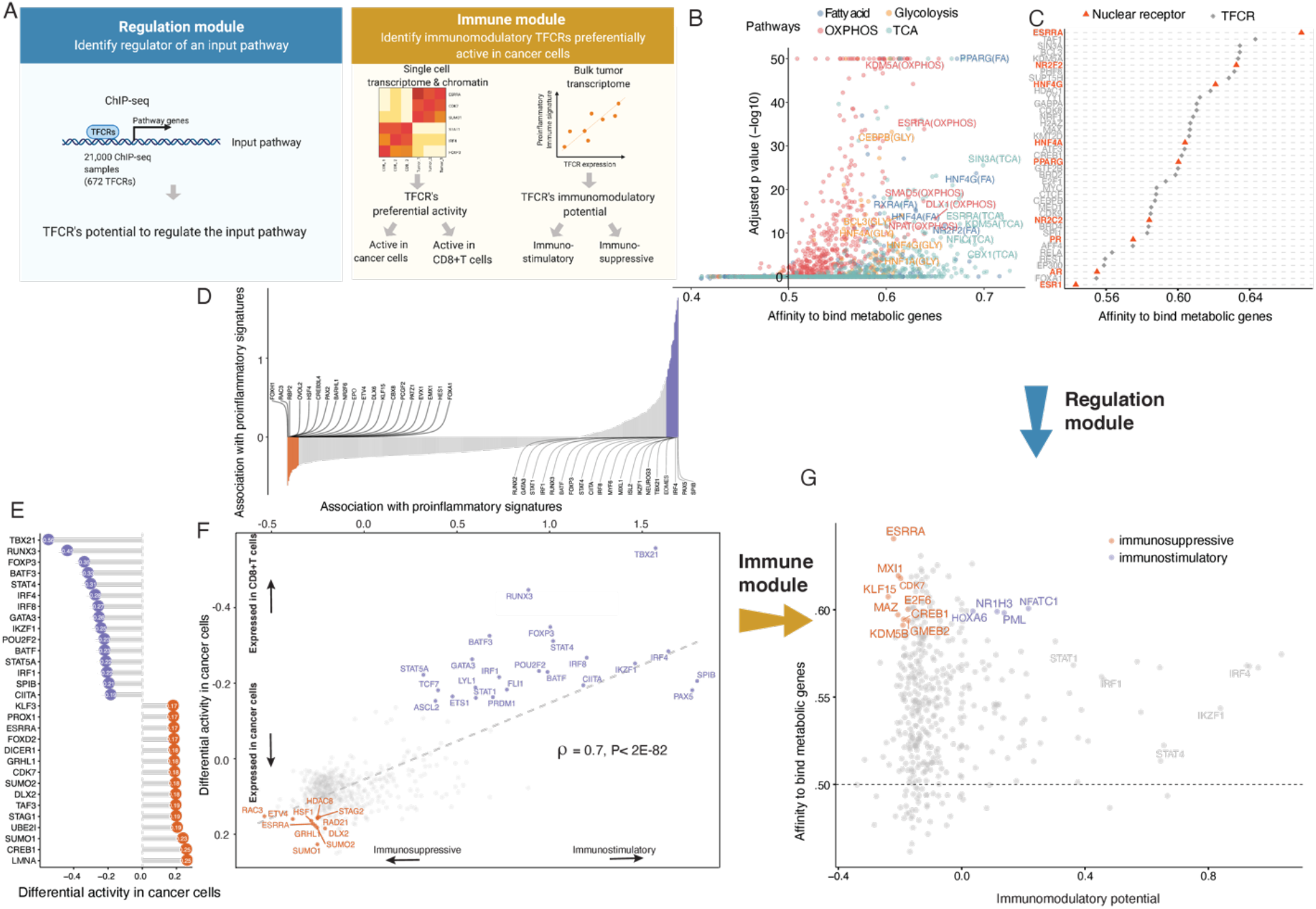
Identification of immune-metabolic regulators: Abbreviations: TFCR, transcription factor and chromatin regulator; RP, regulatory potential of TFCR; TCA, tricarboxylic acid cycle; OXPHOS, oxidative phosphorylation (A) Overall schematic of regulation and immune modules of BipotentR. The regulation module identifies regulators of an input pathway using ChIP-seq data. The immune module distinguishes between putative immunostimulatory vs. immunosuppressive TFCRs using bulk tumor transcriptomes, and identifies TFCRs preferentially active in cancer cells using single-cell tumor transcriptomes. (B) Output of BipotentR regulation module. Affinity and significance of regulators to bind cis-regulatory elements of genes in four energy metabolism pathways. Each dot indicates a regulator, colored by individual pathways. (C) Integrated affinity to bind energy metabolism genes of top predicted master regulators. Nuclear receptors are displayed in red. (D) TFCRs with positive (or negative) associations with proinflammatory signatures are predicted immunostimulators (purple) (or immunosuppressors (orange)). (E) Top TFCRs predicted to be preferentially active in cancer cells (orange) (or CD8^+^T cells (purple)) and their differential activity (estimated from single-cell data). (F) Output of BipotentR immune module: combined association with proinflammatory signatures (D, estimate from bulk RNA-seq) and differential activity in cancer cells (E, estimate from single-cell data) are displayed for each TFCR. (G) Immune-metabolic regulators identified by BipotentR. Energy regulatory potentials (estimated by regulation module) and immune-modulatory potentials (estimated by immune module) of TFCRs. Highlighted TFCRs are significant and among the top 15% in both modules. Immunostimulators (purple) and immunosuppressors (orange) are colored.

BipotentR next identified regulators that can affect multiple energy metabolism pathways (“master regulators”) by ranking TFCRs according to their average overall binding affinities across pathways. Master regulators were enriched in nuclear receptors (P< 1E-7) (Fig 1C), including ESRRA, HNF4A and HNF4G (regulators of liver glycolysis, gluconeogenesis, and FA metabolism (*44*, *45*)) and PPARG (a regulator of adipose tissue energy metabolism (*46*)), along with nuclear hormone receptors such as androgen receptor and estrogen receptor.

Having identified regulators of energy metabolism, we next used the immune module of BipotentR to identify immunomodulatory TFCRs. This module estimates the immunomodulatory potential of ∼700 individual TFCRs from bulk RNA-seq patient tumor data by associating TFCR expression in tumors with levels of a proinflammatory signature. The proinflammatory signature combines 32 key immune response biomarkers, such as mutation burden, neoantigen load, immune infiltration, and interferon-gamma response (*47*) (Methods). TFCR immunomodulatory potential was estimated across several cancer types using a linear mixed model that is robust to cancer-type-specific immune effects (Fig S2A, B) using data from The Cancer Genome Atlas (33 cancer types from 11,000 patients (*47*)). Using two-fold cross-validation, we examined the robustness of immune-potential estimates in cancer-type data held out during prediction. The results (R = 0.99, P < 2.2E-16, Fig S2D) suggest that inferred TFCRs show immunomodulatory properties in several cancer types. TFCRs with the highest inferred immune potential were enriched in immune ontologies, including T-helper differentiation, inflammatory disorders, and viral infection, in addition to carcinogenesis and transcriptional misregulation in cancer (Fig 1D, S2C). Top BipotentR-predicted immunostimulatory TFCRs included well-known regulators of adaptive and innate immunity (e.g., Interferon regulatory factors (*48*, *49*): *IRF1, IRF4, and IRF8*; and the STAT genes *STAT1* and *STAT4* (*48*)), while predicted immunosuppressive TFCRs included regulators of immunotherapy resistance, for example, NR2F6 (*50*). As expected, Predicted immunostimulators were positively co-expressed with immune genes across 5,000 datasets (*43*) (Fig S2E, Methods), while predicted immunosuppressors were negatively co-expressed with immune genes.

The immune module also ensures that suppressing candidate TFCRs specifically block cancer cells without adversely affecting CD8^+^T cells, which are important for anti-tumor immunity (*51*). We achieve this by selecting TFCRs that are active in cancer but not in CD8^+^T cells using single-cell RNA-seq (scRNA-seq) and single-cell ATAC-seq data (scATAC-seq) data. Using scRNA-seq from 5 cancer cohorts (*52*–*56*), TFCRs with significant differential activity in cancer versus CD8^+^T cells across all cohorts are deemed cancer-cell-specific TFCRs (Methods). Using scATAC-seq data (*57*), the module also ensures these TFCRs are functional in cancer cells but not in CD8^+^T cells. To this end, we examined if target genes (inferred by ChIP-seq) of predicted TFCRs are epigenetically accessible in cancer cells (Methods). We found that target accessibility differences between cancer cells and CD8^+^T cells were markedly correlated [Pearson Correlation=.63, P< 2E-59] with scRNA-seq expression differences (Fig S2F), suggesting that predicted TFCRs are active and transcriptionally functional in cancer cells relative to CD8^+^T cells. The top cancer-cell-specific TFCRs included *SUMO1*, *SUMO2*, and *DLX2*, genes known to be tumorigenic and highly active in several cancers (Fig 1E) (*58*–*60*).

We next investigated how these cancer-cell activities (predictions from single-cell data) of TFCRs relate to their immunomodulatory potentials (predictions from bulk tumor data). Strikingly, predicted immunosuppressive TFCRs were preferentially active in cancer cells, while immunostimulatory TFCRs were active in CD8^+^T cells, evident from a strong Pearson correlation of 0.7 [P<2E-82] between immunomodulatory potential and cancer-cell activity (Fig 1F). Thus, inhibiting immunosuppressive TFCRs would be likely to impact cancer cells but less likely to impact CD8^+^T cells adversely. The correlation also shows that prediction from three data types – bulk RNA-seq, scRNA-seq, and scATAC-seq – converged onto a set of immunomodulatory TFCRs, yielding promising targets for anti-tumor immunity. The final set of TFCRs predicted from the immune module were enriched for nuclear receptors (P < 1E-2; Methods), similar to the result from the regulation module.

## Among 38 candidate bipotent immune-metabolic regulator targets, the orphan nuclear receptor ESRRA is most highly ranked

With the two modules developed and validated, we integrated their outputs to identify 38 TFCRs (30 immunosuppressive and 8 immunostimulatory) with immune-metabolic dual functions (Fig 1G, Table S2). Among these were known immune-metabolic TFCRs such as CDK7, which regulates mitochondrial membrane potential (*61*, *62*) and enhances immune suppression (*63*), and NFATC1, which regulates energy consumption and CD8^+^T cell effector function (*64*, *65*).

First, we compared how well genetic inhibition of the 38 identified immune-metabolic regulators suppressed transcription of genes in energy metabolism pathways using a published transcriptome dataset comprised of 570 knockdown/knockout experiments for 308 TFCRs(*66*). We found inhibition of BipotentR-identified regulators markedly suppressed energy metabolism pathways (Fig 2A, P <3.5E-25). Further, inhibition of BipotentR-identified regulators suppressed energy genes more strongly than other TFCRs (Fig S3A, P<7.8E-10), indicating preferential regulation of energy metabolism by the identified TFCRs.

**Figure 2.**
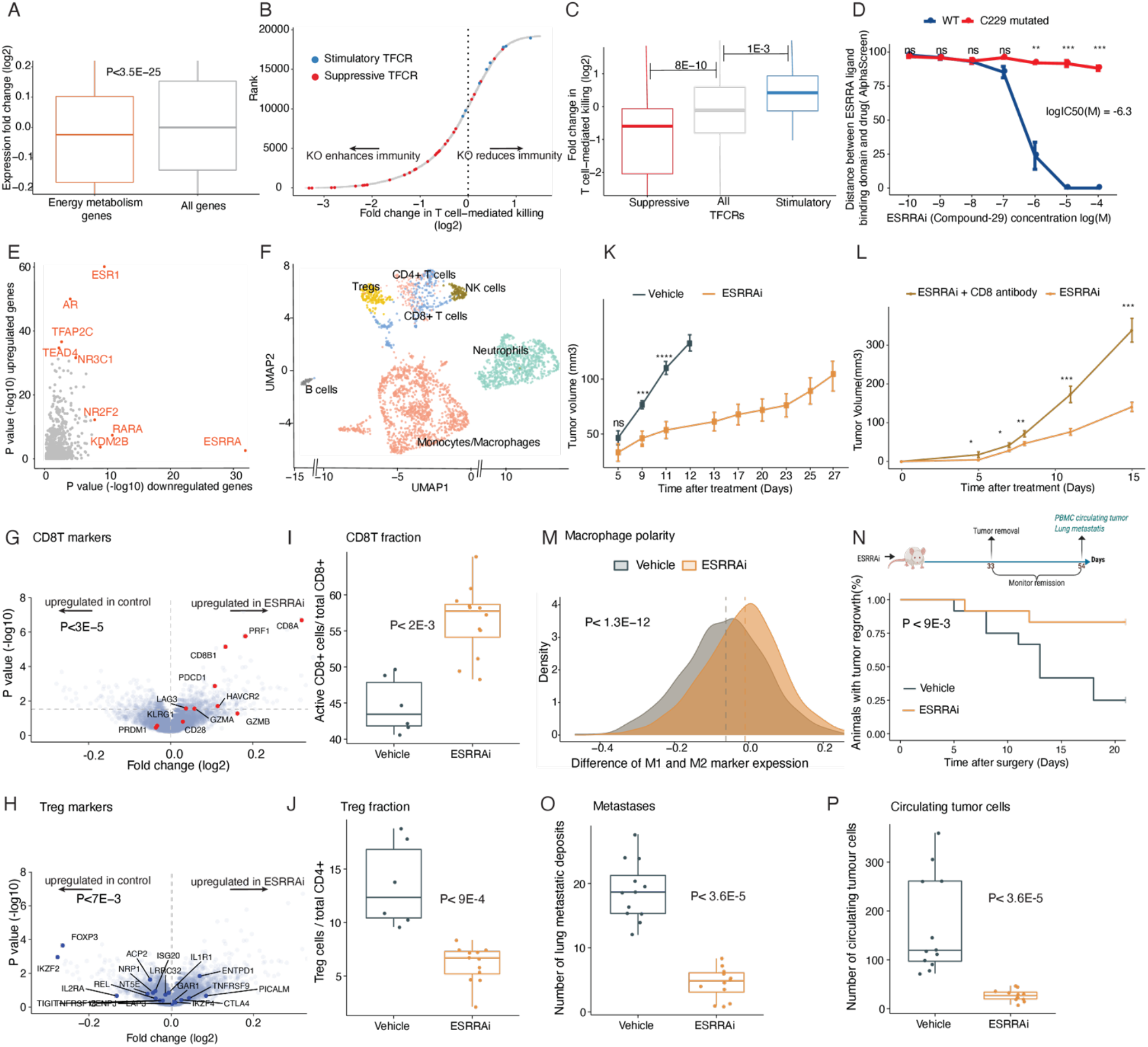
ESRRA inhibition activates anti-tumor immunity in 4T1 mice. P-values using Wilcoxon rank-sum test unless stated otherwise. Tregs, regulatory T-cells; ESRRAi, ESRRA inhibitor. (A-C) Validation of BipotentR-identified bipotent targets. Effect of knockout/knockdown of target identified by BipotentR on (A) expression of genes in energy metabolism pathways, (B, C) T-cell mediated killing of cancer cells. (D) Effect of ESRRAi(Compound-29) concentration on proximity (Alphascreen signal) of the compound to the ligand-binding domain of WT /mutated ESRRA. (E) Transcription regulator analysis (LISA) of up-and down-regulated gene sets by ESRRAi. (F) UMAP display of scRNA-seq of tumor-infiltrating CD45^+^ cells from ESRRAi and vehicle-treated mice. (G, H) Markers of activated CD8^+^T cells (G) and Tregs (H) in genes differential expressed by ESRRAi in lymphoid cells. Significance of up/down-regulation of marker sets estimated using permutation tests. (I, J) Fractions of CD8^+^T cells and Treg cells identified by flow cytometry. (K, L) Tumor volume comparisons between ESRRAi and control (K), and between ESRRAi with and without CD8 antibody (L). (M) Densities of macrophage polarization toward M1 (i.e., for each macrophage cell: average expression of M1 markers - average expression M2 markers), also see Fig S3F, G. (N-P) Measurements done after tumors were surgically removed in ESRRAi or vehicle-treated mice comparing: tumor relapse rate (N), lung metastasis deposits (O), circulating tumor cells in the blood (P).

We next confirmed that inhibition of the identified TFCRs also regulates immunity. We examined a recently published CRISPR screen(*67*) in which cancer cells were subjected to selection by effector T cells to identify gene knockouts that modulate T cell-mediated killing. CRISPR guide RNAs that knockout BipotentR-predicted immunosuppressive TFCRs were depleted (Fig 2B, C; P< 8E-10, n=240 and n=79,481), suggesting that their knockouts enhance T-cell mediated killing. In contrast, guide-RNAs against BipotentR-predicted immunostimulatory TFCRs were enriched, indicating their knockout decreases T-cell mediated killing (Fig 2B, C; P< 1.4E-3, n=64 and n=79,481). Thus, genetic inhibitions of immune-metabolic regulators elicited both immune and metabolic effects.

An orphan nuclear receptor, ESRRA, was predicted to have the highest immune-metabolic potential. Targeting ESRRA in immuno-deficient models has been shown to inhibit tumors by direct cell-intrinsic mechanisms (*37*, *68*), and we hypothesized that targeting ESRRA would also inhibit tumors by immune-mediated mechanisms. To illustrate a proof-of-principle bipotent target, we set out to determine the dual potential and clinical relevance of ESRRA in different cancer types.

## Inhibition of ESRRA (ESRRAi) stimulates anti-tumor immunity

We evaluated inhibiting ESRRA by small interfering RNA (siRNA) and two structurally similar diaryl ether based thiazolidinediones, which function as selective ligands against ESRRA (Compounds 29 and 39 from Johnson and Johnson (*69*)). We first tested both small-molecule inhibitors *in vitro* by mutating the known compound binding site in the ESRRA ligand-binding domain (C229 site) and showing that the mutation rescued the ESRRA inhibition (Fig 2D, S3B). Next, we investigated siRNA and compound-39 for on-target and off-target effects through RNA-seq (Methods). Both approaches selectively suppressed putative ESRRA genes targets (that were identified from ESRRA ChIP-seq data) (Fig S3C). An unbiased prediction of 700 putative regulators of the genes differentially expressed upon the two approaches (*70*) yielded ESRRA as the top regulator of down-regulated genes (topmost for drug inhibition, Fig 2E; second-highest for siRNA, Fig S3D). These analyses showed both inhibitions selectively suppress ESRRA and have limited off-target effects. We chose to pursue drug inhibition of ESRRA (“ESRRAi”, which refers to inhibition by Compound 29 or Compound 39) because of its translational potential and somewhat superior potency in targeting ESRRA. Compound-29 is known to be more stable metabolically in human microsomes than compound-39 (*69*), and therefore we used compound-29 for *in vivo* testing (Methods)..

We next tested if ESRRAi could induce anti-tumor immunity in two immunosuppressive murine tumor models: 4T1 (triple-negative breast cancer) and B16F10 (melanoma). We treated the 4T1 mice with ESRRAi or vehicle control and surgically resected their tumors. We performed scRNA-seq of CD45^+^ cells sorted from tumors, clustered and annotated cells using classical markers, and identified major tumor-infiltrating immune cells in both conditions (Fig 2F, Methods). We initially studied immune cells of lymphoid lineage for changes in their fraction by ESRRAi treatment and found higher CD8^+^T cell infiltration with the treatment (Fig S3E). A CD8^+^T cell marker, *Cd8a*, was the topmost upregulated gene in the lymphoid lineage of ESRRAi-treated tumors compared to controls (Fig 2G). Markers of activated CD8^+^T cell (Fig 2G), including perforin and granzymes, were also upregulated [P<3E-5], suggesting that infiltrating CD8^+^T cells in treated tumors were also activated. We also showed increased infiltration of activated CD8^+^T cells with ESRRAi in tumors [P<2E-3] using fluorescence-activated single-cell sorting (FACS) (Fig 2I).

Next, we asked if ESRRAi-induced infiltration of activated CD8^+^T cells exerts an anti-tumor effect. ESRRAi treatment markedly reduced tumor growth (Fig 2K). Two lines of evidence linked this tumor elimination with CD8^+^T cells. First, among ESRRAi-treated mice, those with higher CD8^+^T infiltration showed superior tumor elimination [Spearman correlation = −0.62](Fig S3I). Second, CD8^+^T cell depletion abrogated the anti-tumor effect of ESRRAi (Fig 2L). Another ESRRAi-induced change in the lymphoid lineage was downregulated markers of regulatory T cells (Tregs) (Fig 2H). Correspondingly, lower Treg infiltration in the ESRRAi condition was observed in single-cell data (Fig S3E), which was further confirmed using FACS [P<9E-4] (Fig 2J), indicating that ESRRAi treatment suppressed Treg infiltration into tumors. These analyses revealed the specific roles of different T-cell populations in ESRRAi anti-tumor immunity.

ESRRA deficient mice in a non-cancer context have shown macrophage-mediated inflammation (*71*). Therefore, we postulated that ESRRAi might affect tumor macrophages. Indeed, monocytes/macrophages were polarized toward proinflammatory M1 (Fig 2M, S3F) in the ESRRAi-treated tumors. In contrast, macrophages were polarized toward pro-tumorigenic M2 in controls (Fig S3G). Moreover, monocytes/macrophages of treated tumor expressed M1 markers (*Tnf, Ccl5, Nos2, and Il1a*) (*72*) and downregulated M2 markers (Fig S3H).

Next, we tested the effect of ESRRAi treatment on tumor relapse from minimal residual disease. After surgically removing 4T1 tumors (Fig 2N), ESRRAi treated mice experienced significantly fewer tumor relapses (Fig 2N); moreover, their relapsed tumors had significantly attenuated growth (Fig S3J). We examined incised lungs from treated mice and observed fewer lung metastatic deposits than the control group (Fig 2O). We also cultured the circulating tumor cells from the blood of treated mice and observed a significant decrease in the number of colonies relative to the control group (Fig 2P). These data suggest that ESRRAi can prevent relapse of surgically resected tumors.

Similar ESRRAi anti-tumor responses were observed in an additional immune-cold tumor B16F10 mouse model and two formulations (Solutol and PEG) (Fig S4, Methods). Thus, our data indicate that ESRRAi induces anti-tumor effects that depend on T-cells and polarizes macrophages toward a more proinflammatory state.

## Immune signaling pathways link ESRRAi to immune response

We next asked what cell-autonomous immune-metabolic pathways underlie ESRRAi anti-tumor immunity. We treated a human breast cancer cell line (SKBR3) with ESRRAi and measured transcriptomic changes at three timepoints. ESRRAi suppressed metabolic genes at all time points, particularly energy metabolic pathway genes (Fig 3A, B; S5A). ESRRA suppressed using siRNA also inhibited energy metabolism pathways (Fig S5B, C), confirming this observation.

**Figure 3.**
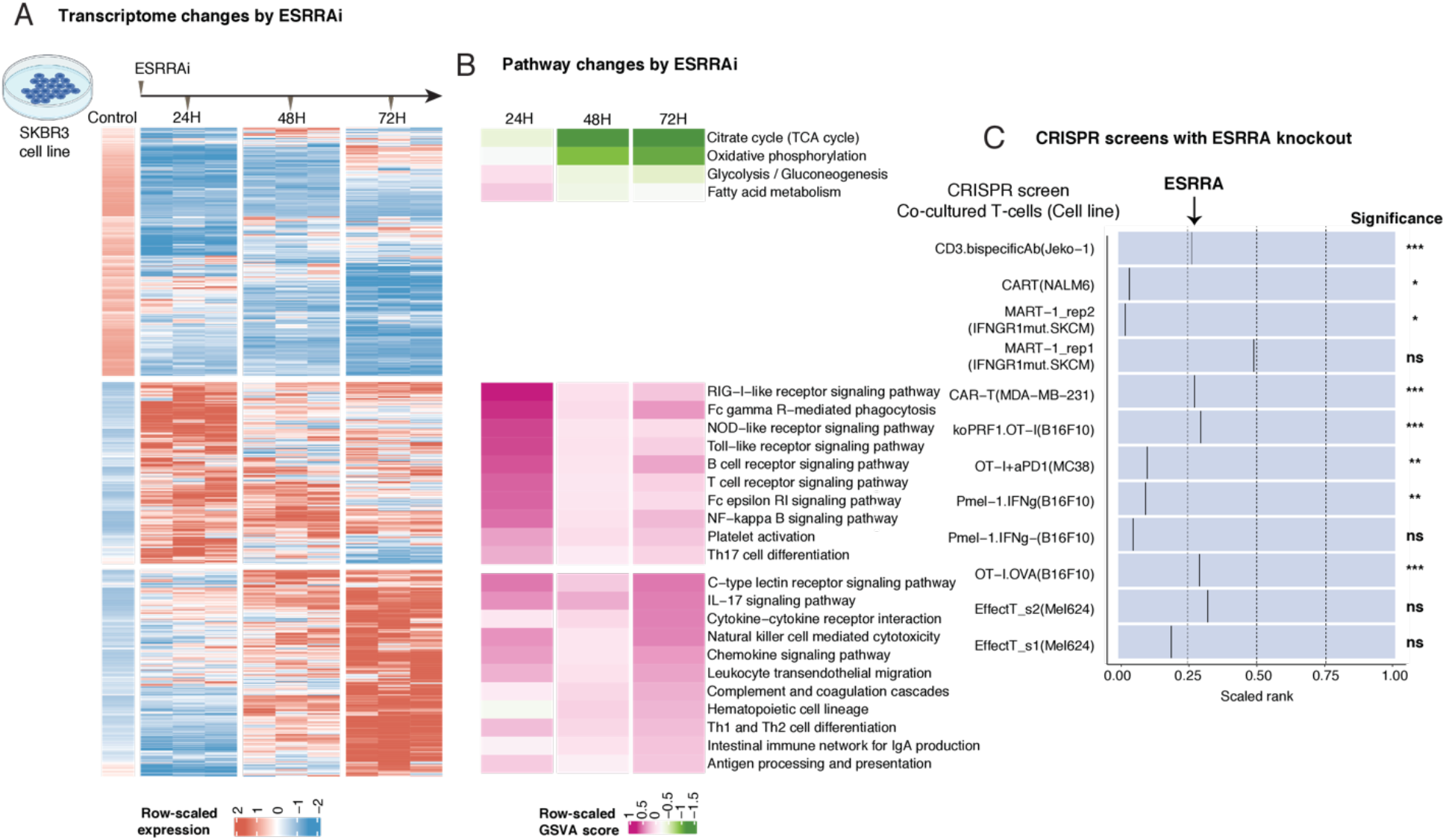
Signaling induced by ESRRAi *in vitro*. Abbreviations: GSVA, Gene Set Variation Analysis (A) Differentially expressed genes between ESRRAi and control in the SKBR3 cell line at three time points. Genes were clustered by K-means. (B) Pathway enrichment scores corresponding to clusters of differentially expressed genes shown in (A). (C) ESRRA knockout potentiates T-cell killing as observed in CRISPR knockout screens in cancer cells co-cultured with T-cells. The black line represents the relative position of ESRRA knockout among all gene knockouts ranked from most depleted to least depleted. The significance of ESRRA knockout in screens is also displayed.

In contrast, the effect on immune pathways showed a striking temporal trend: the treatment upregulated innate immune signaling at 24h, while at 72h, it upregulated adaptive immune signaling (Fig 3A, B). Twenty-four hours after the treatment, the treatment upregulated signaling of toll-like receptor (TLR), which is known to stimulate antigen presentation(*73*). At this time point, the treatment also upregulated signaling of Fc-epsilon-RI, Rig-I-like, and NOD–like receptors, which along with TLR signaling, are known to promote inflammatory cytokine secretion(*74*). Accordingly, the ESRRAi treatment upregulated genes involved in antigen presentation (Fig S5D; 3B) and cytokine-cytokine receptor interactions – especially cytokines that polarize macrophages towards M1 (Fig S5E; 3B) – at the 72h time point. This ESRRAi-induced upregulation of macrophage-polarizing cytokines is consistent with macrophage polarization by ESRRAi observed in our *in vivo* single-cell experiments (Fig 2M). The ESRRAi treatment also upregulated 20 immunomodulatory TFCRs identified by BipotentR (Fig S5F, Enrichment P < 4.7E-12), suggesting that ESRRA is an upstream regulator of other immune regulators.

We next examined if the knockout of ESRRA in cancer cells induces antigen presentation by analyzing data from CRISPR knockout screens (*75*–*78*) designed to identify regulators of type-I antigen presentation genes (MHC-I). These screens sort cancer cells transduced with gRNA into low or high MHC-I groups based on their MHC-I protein expression (Fig S6E). gRNAs that knockout (KO) ESRRA were enriched in high MHC-I and depleted in low MHC-I groups (Fig S6E), confirming that ESRRA knockout increases MHC-I antigen presentation.

Since increased MHC-I antigen presentation in tumors enhances the ability of T-cells to kill cancer cells (*67*, *79*), we hypothesized that ESRRAi would enhance tumor killing by T cells. We tested this using published CRISPR screens that co-culture cancer cells with T-cells to identify which gene knockouts in cancer cells enhance their T-cell–mediated killing (*67*, *80*–*83*). ESRRA KO potentiated the killing of cancer cells by both patient-derived and engineered effector T-cells in various experimental and cell-line contexts (Fig 3C). Because T-cell-mediated killing has previously been shown to be enhanced by OXPHOS suppression (*27*, *67*), we asked if OXPHOS targets of ESRRA (derived from ESRRA ChIP-seq (Methods)) can explain this effect. Indeed, knockout of *COX10*, *ATP51B*, *NDUFA6* alone not only potentiated T-cell-mediated killing (Table S3), but also increased protein levels of antigen presentation genes (Table S4). Thus, OXPHOS suppression by ESRRAi can explain the activation of antigen presentation and T-cell-mediated immunity by ESRRAi.

Finally, we generated a signature based on differential expression upon ESRRAi. Using this signature, we divided the cell lines in the 1,000 Cancer Cell Line Encyclopedia data (*84*) by high and low ESRRA activity. ESRRA activity was correlated with cell-autonomous immune-metabolic effects broadly across cancer types represented in the Cancer Cell Line Encyclopedia, such that cell lines with low ESRRA activity exhibited decreased energy metabolism and upregulated immune pathways (Fig S6A-D; see Supplementary Text).

## ESRRA activity in patient tumors correlates with antigen presentation, immune cell infiltration, and macrophage polarization

Unlike experimental screens, BipotentR derives immune-metabolic targets directly from patient data, which captures the patient tumor immune microenvironment, increasing the potential clinical relevance for candidate targets. To evaluate the potential clinical relevance of ESRRA, we compiled an additional set of patient tumor transcriptomes from more than 200 bulk and 78 single-cell cohorts and investigated the correlation between ESRRA activity and the immune effects that were observed *in vivo* after ESRRAi treatment.

First, we analyzed 33,000 tumor transcriptomes compiled from several cohorts (*85*), including TCGA and PRECOG datasets (*86*). ESRRA activity in tumors was quantified as the weighted sum of expression of ESRRA targets (derived using gene signatures induced by ESRRA inhibition (Methods)). In tumors with low ESRRA activity, tumor energy metabolism was significantly downregulated across several cancer types in TCGA (Fig S7A; Methods). In these tumors, cytokine interaction pathways were upregulated (Fig S7B), consistent with the *in vitro* induction of macrophage-polarizing cytokines upon ESRRAi. We, therefore, asked whether macrophage polarity was also shifted in such tumors. We analyzed how macrophage polarity (estimated using gene expression signature (*87*)) relates to ESRRA activity in tumors, and found that macrophage polarity was markedly correlated with ESRRA activity within tumors across most cancer types (Fig 4A), strongly suggesting that M1 macrophage polarization upon ESRRAi seen *in vivo* in mice may be clinically relevant in most cancer types.

**Figure 4.**
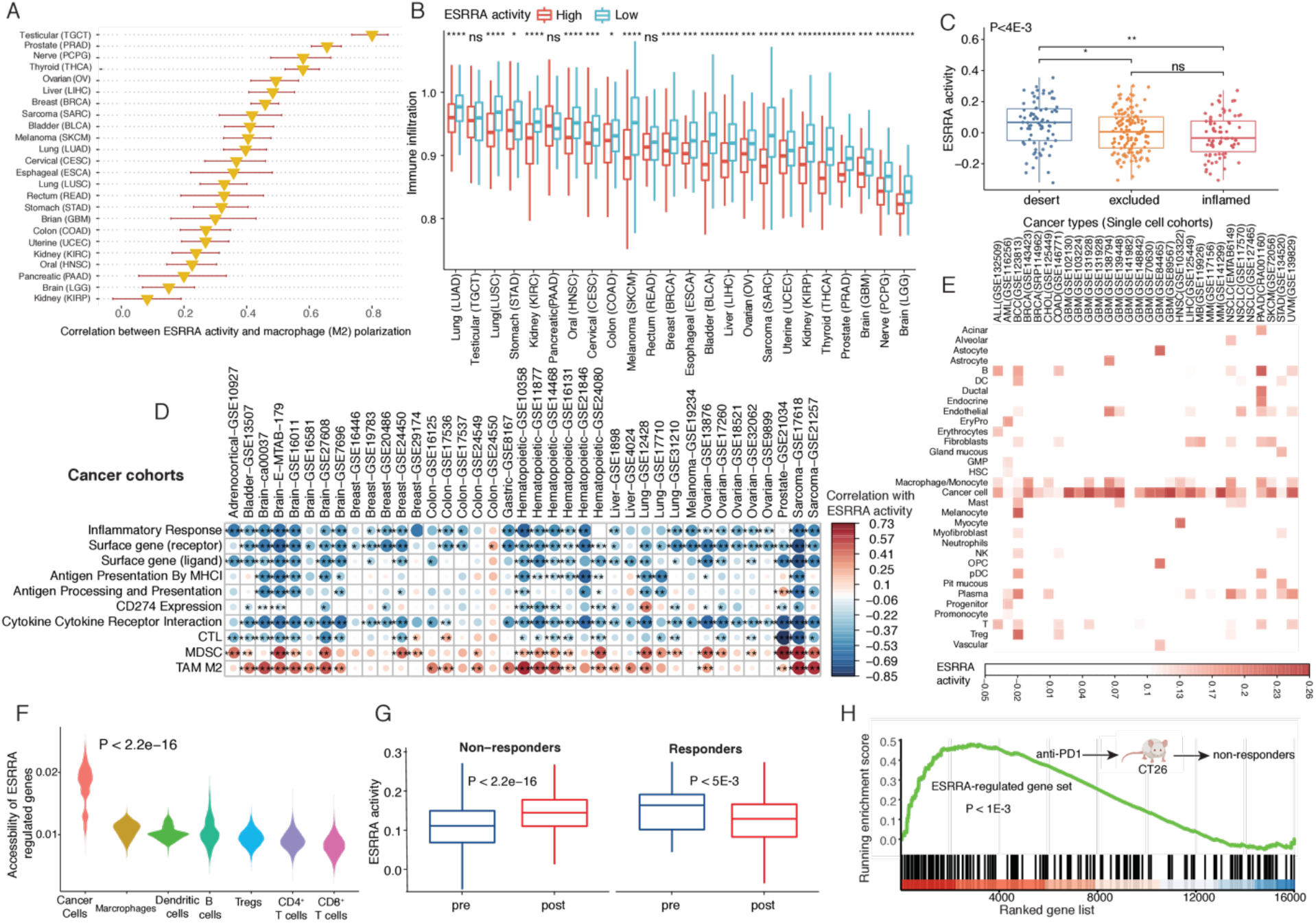
Clinical significance of ESRRAi. CTL, cytotoxic T cell; MDSC, Myeloid-derived suppressor cells; TAM, Tumor-associated macrophages. P-values estimated by Wilcoxon rank-sum test. MHC, major histocompatibility complex (A) Correlation of ESRRA activity with macrophage polarization towards M2 across TCGA cancer types. Correlation coefficient and standard error are displayed (B) Immune infiltration in low and high ESRRA activity tumors across cancer types in TCGA. (C) ESRRA activity in patient tumors with inflamed, excluded, and desert immunophenotypes ( based on CD8^+^T infiltration levels: inflamed > excluded > desert) in bladder cancer cohort. P-value was estimated using Kruskal-Wallis test. (D) Correlation between ESRRA activity and immune biomarkers in PRECOG collection of patient cohorts. (E) ESRRA activity in cancer and immune cells of 30 scRNA-seq cohorts. (F) The chromatin accessibility of ESRRA targets in different cell types from scATAC-seq data of a skin cancer cohort. (G) Cancer cell ESRRA activity in patient (skin cancer) tumors pre- and post- anti-PD1 treatment for responders and non-responders. (H) Enrichment analysis of ESRRA-regulated gene set in non-responding CT26 mice after anti-PD1 treatment.

Antigen presentation promotes immune infiltration into tumors. Antigen presentation genes were upregulated in tumors with low ESRRA activity across several cancer types (Fig S7C). If antigen presentation simulated by ESRRAi is clinically relevant, tumors deficient in ESRRA activity should have elevated immune infiltration. Indeed, tumors with low ESRRA expression had high immune infiltrations, including CD8^+^T cell infiltration, across most cancer types (Fig 4B, P < 2E-16 controlled for cancer types, Fig S7D). We evaluated high CD8^+^T infiltration in low-ESRRA activity tumors in a cohort of more than 300 bladder cancer patients where tumor CD8^+^T cell infiltration was measured by immunohistochemistry (*88*). Tumors with the highest levels of CD8^+^T cell infiltration (Immune-inflamed tumors) had the lowest ESRRA activity, followed by immune-excluded tumors (which have CD8^+^T cell infiltration, but not proximal to tumor cells) with intermediate-levels of ESRRA activity, while CD8^+^T cell deficient tumors showed the highest ESRRA activity (Fig 4C; Fig S8A, B). Dramatic upregulation of M1-polarizing cytokines and antigen presentation genes (Fig S8C, D) were also seen in tumors with low ESRRA activity from this cohort (see Supplementary Text). Finally, we found that low ESRRA activity in tumors is associated positively with proinflammatory factors, and negatively with anti-inflammatory factors in both TCGA (Fig S8E) and PRECOG data (Fig 4D). Together, the patient tumor data demonstrate the potential clinical relevance of ESRRAi in enhancing antigen presentation, immune cell infiltration, and macrophage polarization in multiple cancer cohorts and cancer types.

## ESRRA activation in immunotherapy-resistant tumors

Targeting tumor energy metabolism by ESRRAi would be detrimental if it also affects T-cell metabolism (*27*, *89*). To validate BipotentR’s prediction of cancer-cell specificity for ESRRA inhibition and to further investigate the effect of ESRRAi on T-cells and cancer cells, we compiled and analyzed 78 single-cell transcriptome datasets from patients with 27 different major cancer types (*90*) (Methods). We found that ESRRA was expressed at the highest levels in cancer cells, but that it was also expressed at lower levels in macrophages and T-cells (Fig S9A). Since the functional activity of a nuclear receptor depends not only on its expression but also on its ligands, cofactors, and stimulation, we reasoned that despite being expressed in T cells, ESRRA might have low functional activity in T cells. Indeed, ESRRA activity levels, quantified as the expression of ESRRA targets, were lowest in T cells (Fig 4E, Methods). In contrast, the highest and second-highest levels of ESRRA activity were observed in cancer cells and M2 macrophages (Fig 4E, S9B-D).

Next, we examined ESRRA cell-specific function by comparing accessibility of its target genes (inferred from ESRRA ChIP-seq, Methods) in different cell types using scATAC-seq data from non-melanoma skin cancer patients (*57*). ESRRA target gene accessibility was highest in cancer cells, second highest in macrophages, and lowest in CD8^+^T cells (Fig 4F), consistent with ESRRA activity distribution in scRNA datasets. The data support a model in which ESRRA has a higher level of activity in cancer cells relative to CD8^+^T cells as measured by gene expression, target transcription, and chromatin accessibility. Thus, ESRRAi likely has a lower impact on the energy metabolism of CD8^+^T cells.

To specifically test whether ESRRAi treatment impacts energy metabolism of CD8^+^T cells, we analyzed scRNA data from CD45^+^ cells from our *in vivo* 4T1 mouse model, in which the anti-tumor effect of ESRRAi was clearly measurable. ESRRA activity in CD8^+^T cells was unchanged in ESRRAi treated mice relative to control (Fig S10A), whereas macrophages/monocytes were the only CD45^+^ cells that showed a decrease in ESRRA activity post-treatment. We also evaluated our scRNA data using Augur (*91*), a method that identifies cell types affected by treatments, which also found no significant cell-intrinsic changes in CD8^+^T cells post-ESRRAi (Fig S10B). ESRRAi treatment in 4T1 mice did not significantly change body weights or health parameters (Fig S10C), which we further confirmed in the B16F10 mouse model (Fig S10D), suggesting that ESRRAi treatment was not nonspecifically toxic. These data strongly suggest that ESRRAi has a little adverse impact on CD8^+^T cells.

Next, we studied the effect of immunotherapy on ESRRA activity. Analysis of a cohort of immunotherapy-resistant melanoma patients (*92*) revealed an intriguing trend: cancer cells from post-immunotherapy tumors had markedly higher ESRRA activity than those from pretreatment tumors (Fig S10E). As the cohort only contained immunotherapy-resistant patients, we asked whether the trend is specific to resistant patients or is also present in responders. To that end, we analyzed a non-melanoma skin cancer scRNA-seq cohort (*93*) containing both immunotherapy responder and resistant patients. Indeed, the trend of immunotherapy-induced ESRRA activity increase was specific to immunotherapy-resistant tumors (Fig 4G). In fact, in responders, ESRRA activity decreased in cancer cells upon immunotherapy (Fig 4G). This data is consistent with the hypothesis that immunotherapy-resistant tumors achieve high levels of immune suppression via ESRRA.

To experimentally test the trend of increased ESRRA activity and resulting immune-suppression upon immunotherapy, we chose a syngeneic mouse model of colorectal cancer (CT26) known for its heterogeneous immunotherapy (anti-PD1) response (*94*). CT26 mice were treated with anti-PD1, and bulk tumor RNA-seq was conducted in responding and immunotherapy-resistant mice to assess treatment effects on ESRRA target genes. Target genes of ESRRA were enriched in genes upregulated in immunotherapy-resistant mice (Fig 4H) but not in responders (Fig S10F), suggesting that ESRRA activity increases in immunotherapy-resistant tumors post-anti-PD1 treatment. Increased ESRRA activity was also correlated with decreased CD8^+^T infiltration and increased M2 macrophages in tumors (Fig S10G).

This *in vivo* experiment supports a model in which immune checkpoint blockade increases ESRRA activity, specifically in immunotherapy-resistant tumors. While increased ESRRA activity elevates immune suppression, it may also increase the vulnerability of immunotherapy-resistant tumors to ESRRA inhibition. The potential vulnerability is also supported by our *in vivo* experiment showing ESRRAi effectiveness in 4T1 and B16F10 models – both of which respond poorly to ICB. Future clinical investigations are required to reveal whether immunotherapy-resistant tumors benefit from the immunostimulatory effect of ESRRA inhibition. Overall, our studies show targeting ESRRA induces proinflammatory cytokines, which in turn polarize macrophages toward proinflammatory states. Inhibition of ESRRA by CRISPR or drugs stimulates antigen presentation genes, which in turn recruit effector CD8^+^T cells to tumors and enhance tumor elimination by T-cells (Fig S10H).

## Application of BipotentR to other pathways

We also identified 14 bipotent TFCRs that simultaneously regulate angiogenesis and immune response (Table S5) and 14 TFCRs that regulate evasion of growth suppressors and immune response (Table S6), using these pathways as inputs to BipotentR (Methods). Using existing CRISPR datasets (*67*), we tested if genetic inhibitions of these 28 identified TFCRs can elicit dual anti-tumor efficacy. CRISPR knockouts of these TFCRs markedly improved the killing of cancer cells by T-cells (Fig S11A, P<1.3E-6). Knockdown or knockout of the identified TFCRs suppressed genes involved in angiogenesis or evasion of growth suppressors (Fig S11B, P<8.1E-24), and the suppressed genes were preferentially regulated by the bipotent TFCRs (Fig S11C, P< 9E-10).

## Conclusion

We show that BipotentR identified ESRRA as a candidate target for a bipotent drug that affects both tumor immunity and energy metabolism. ESRRA inhibition affected energy metabolism, including OXPHOS, in cancer cells without curtailing CD8^+^T cell activity. In cancers, targeting ESRRA induced cytokines, which in turn polarized macrophages toward proinflammatory states. CRISPR or drug targeting of ESRRA stimulates antigen presentation genes (Fig S6E, S5D), which in turn recruits effector CD8^+^T cells to tumors (Fig 2G, I) and enhances tumor elimination by T-cells (Fig 2K, L, and 3C). These findings are strongly supported by recent studies showing that OXPHOS suppression can alter both T cells (*27*, *67*) and macrophages (*95*–*97*). High ESRRA activity was observed in immunosuppressive and immunoresistant tumors across several cancer types (Fig 4A-D), strongly suggesting clinical relevance in patient tumors. We show that BipotentR is generalizable to other biological pathways. These data demonstrate the advantages of suppressing energy metabolism in selected cell types while simultaneously stimulating an immune response with a single drug and provide proof-of-concept for BipotentR in bipotent drug discovery.

## Code, website, and data availability

The BipotentR is available at http://bipotentr.dfci.harvard.edu/. The R-package BipotentR is available at (https://github.com/vinash85/TRIM). Upon publication, a user-friendly website will be created that will identify bipotent regulators that simultaneously modulate immune response and any given input pathway or pathway list. Bulk and single RNA-seq data generated from the current study are available at (https://www.dropbox.com/sh/qoqlx9724k0k869/AADZe7oZp0vs4gKzqeRwTUjaa?dl=0) and will be submitted to a public repository upon publication.

